# Diverse Patterns of Allele-Specific Expression in Healthy Human Tissues

**DOI:** 10.1101/2025.10.14.682127

**Authors:** Blaise L. Mariner, Bryan Sands, Soo Yun, Tiffany Jones, Elizabeth M. Swisher, Mark A. McCormick, Alexander R. Mendenhall

## Abstract

Differences in gene sequence and gene expression underlie variation in traits. However, even monozygotic twins do not express their genes in the same way, develop divergence in traits, and succumb to distinct chronic diseases. During development, epigenetic silencing programs cause diversity in allele expression, resulting in differences in traits and chronic disease risk. To quantify human autosomal allele expression between individuals, we analyzed human allele-specific expression data from the GTEx project. For hundreds of genes, some individuals will express the gene biallelically, while many others may only express one allele or extreme bias towards one allele. We found gene-specific patterns of interindividual variation in allele bias. We found that some individuals have more genome-wide monoallelic/biased expression than others. Individuals also had distinct combinations of allele expression bias. These differences can underlie variation in traits, idiopathic or incompletely penetrant traits/diseases, and chronic diseases.

**Significance/Impact:** Allele-specific expression can affect cancer, immune response, and genetic disease. This work reveals 1) gene-specific multimodal patterns of interindividual variation in allele bias, 2) that individuals can maintain bias across tissues, and 3) that different individuals have distinct combinations of silenced alleles. These different patterns and the weighted classifications demonstrate how allele bias manifests between individuals; there are individuals and tissues with more biased/non-Mendelian expression and some tissues have more age-related changes in which alleles are silenced.

## Introduction

Differences in genes and *external* environments do not explain all the variations in biological traits/outputs. Sequence variation can influence organism phenotype by causing changes in the quality or quantity of gene expression (Buckland 2004). This can occur in both protein-coding and non-coding regions and impact genetic disease (Antonarakis et al. 1984; Koivisto et al. 1994). Yet, despite over a century of standardization of laboratory conditions and inbreeding to minimize biological variation in traits, isogenic laboratory animals are never exactly alike. The same is true for monozygotic human twins who live for different amounts of time and succumb to distinct diseases. Genes and environments together can explain all variation in traits in part because the term environment actually includes intrinsic variation that is not attributable to the *external* environment. There are intrinsic programs that generate physiological differences, even among genetically identical individuals, by randomly/probabilistically silencing alleles of thousands of autosomal alleles during early development (Chess 2016).

Random autosomal monoallelic expression has been linked to differences in genetic and chronic diseases (C. Lee et al. 2019; Luft et al. 2020; Izzi et al. 2016; McKean et al. 2016; Falkenberg et al. 2017; Castel et al. 2018; Bielski et al. 2018; Huang et al. 2015; Adegbola et al. 2015). People can succumb to recessive genetic diseases despite having only one recessive allele when they silence the non-pathological allele. People can also escape genetic disease by silencing a dominant pathological allele. For example, in a case report from Japan, only patients that expressed the dominant *PIT1/POU1F1* allele got pituitary disease, despite several members of the family carrying the allele in a silenced state. (Okamoto et al. 1994). Allele expression bias in some specific contexts may aid organisms in bypassing neoplastic progression (Yan et al. 2002). Allele expression bias for certain genes may function as a risk factor for cancer development and the onset of other genetic diseases (Castel et al. 2018). Recent work has also shown that monoallelic expression affects the penetrance of immune traits in human T cells (Stewart et al. 2025). Finally, recent work also analyzing GTEx data found that genes expressed in a biased/monoallelic fashion are enriched for effects on aging and chronic disease (Kravitz et al. 2023).

At the molecular level, allele preference is affected by cis and trans factors that can place people into multimodal classes of gene expression caused by different states of allele silencing. Recent studies reveal that that introns can affect the degree of allele bias in *Caenorhabditis elegans* (Sands, Yun, and Mendenhall 2021). MAE can be affected by DNA methylation (Gupta et al. 2022) and H3K9 histone methylation (Sands et al. 2024; Eckersley-Maslin et al. 2014) at promoters, as well as by H3K27 and H3K36 histone methylation in the gene body (Stewart et al. 2025; Nag et al. 2015, 2013). Random autosomal monoallelic expression can span the full spectrum of bias, from completely silenced to completely biallelic (Raser and O’shea 2004; Sands Yun and Mendenhall 2021). That is, for some genes, model systems indicate that there will be individuals who have tissues with biallelic expression, and other individuals who have monoallelic or very biased expression of that exact same gene in the exact same tissue. Work in genetic model organisms has shown that this random bias is set during early development and does not appear to be heritable (Sands et al. 2024).

Here, we show strikingly distinct inter-individual allele expression patterns as scatter plots generated by analyzing the haplotype expression of open reading frames (ORFs) from the human GTEx v8 haplotype expression dataset, which contains data from 54 human tissues from 838 individuals (**Figure 1A**) (THE GTEX CONSORTIUM 2020; Castel et al. 2020). We used a conservative approach to classifying genes that showed relatively more or less biased expression patterns across tissues using the GTEx data. We used unsupervised machine learning to cluster 17,406 ORFs less susceptible to read noise by requiring that ORFs have adequate reads across samples (See Methods). We then used unsupervised machine learning-based hierarchical clustering of these ORFs, which identified four main types of genes: biallelic, slightly biased, very biased, and monoallelic. This approach is distinct from other complementary approaches classifying monoallelic or randomly silenced/biased genes in the human genome (Nag et al. 2015; Kravitz et al. 2023; Savova et al. 2016). Our specific results are shown below.

**Figure 1:**
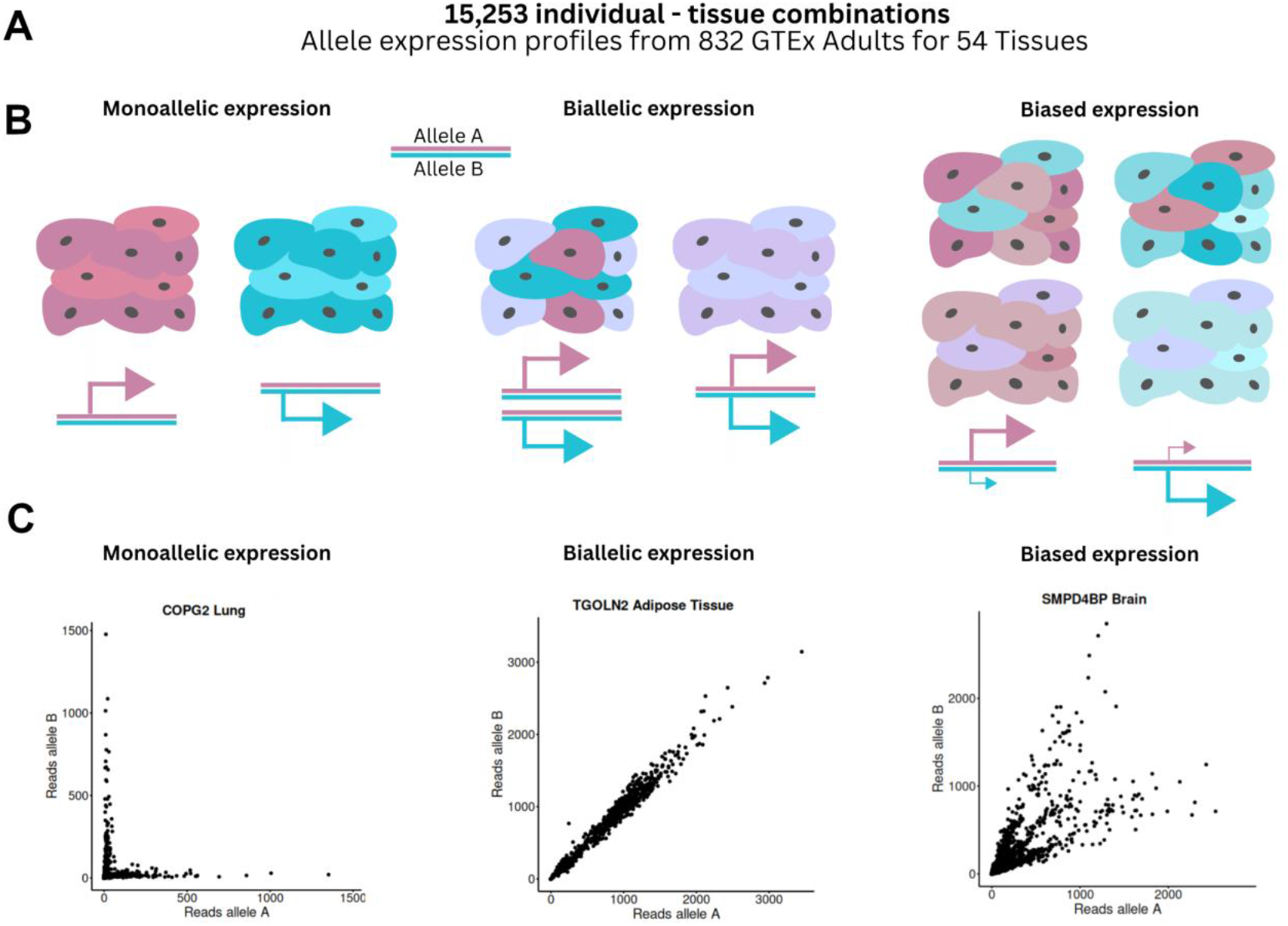
GTEx RNAseq data can show the degree of organs’ allele preference. A) The GTEx v8 bulk RNAseq dataset comprises 54 tissue types and 864 individuals. This bulk RNAseq dataset can capture tissues’ preference toward one allele or another. It cannot capture variegated allele-specific expression/silencing that varies in a cell-autonomous manner in a tissue - a patchwork - that comes out as biallelic with bulk tissue RNA-seq, but not with bulk Chip-seq or Nanopore sequencing - both of those technologies would show 50-75% on versus 50-25% off, in a tissue that was variegated/patchwork. The present study identifies genes with bias in allele expression that is propagated throughout a tissue. B) Cartoon examples of what allele bias propagated in a tissue can look like at the cellular level with differently colored alleles. C) Scatter plots of human allele expression data from different tissues show distinct monomodal, bimodal, trimodal or continuous patterns of gene expression when considering both alleles.

## Results

### Scatter Plots Show Distinct Patterns of Inter-individual Variation in Allele Expression Bias

For every gene in every tissue in the GTEx dataset, we were interested in determining how inter-individual variation in allele expression manifested. We used scatter plots, which have proven useful for understanding cell-to-cell variation in allele expression (Sands, Yun, and Mendenhall 2021; Gupta et al. 2022; Burnaevskiy et al. 2019),. The GTEx dataset consists of bulk sequencing data, not single-cell data, so we plotted a person’s average allele bias for a tissue. We previously observed biallelic, biased, and monoallelic expression patterns in whole tissues in animal models and cell culture (Sands, Yun, and Mendenhall 2021; Gupta et al. 2022; Burnaevskiy et al. 2019; Sands et al. 2024), so we expected to find those patterns. We plotted the reads from each gene allele in each tissue in each person for samples that met our minimum quality standards (Methods).

We found that scatter plots of allele read from the GTEX data revealed striking patterns of inter-individual variation in allele expression bias, shown in **Figure 1**. We saw several patterns of allele bias. Some genes are biallelic, some are biased, and some are completely monoallelic. We found biallelic genes that all individuals would express both alleles equally, which showed the expected “cone” patterns. But, in some biallelic expression cases, genes did not have the expected maintained coefficient of variation at higher expression levels and instead decreased the relative deviation between alleles, revealing “stretched ellipse” patterns of interindividual variation in allele expression bias. Other genes showed patterns of interindividual differences in allele expression, indicating that individuals’ tissues could be in monoallelic or biased states of gene expression. For some genes in some tissues, individuals are found to have monoallelic expression of one allele or the other, forming a bimodal “L” pattern. Other genes showed striking, three-state/trimodal/chicken foot patterns, wherein individuals’ tissues expressed alleles exclusively in three states: very biased/monoallelic or biallelic, with a clear separation between the three states. Finally, there were also instances of a continuous distribution of allele expression bias states across the population, with those scatter plots revealing “fan” patterns. We next sought to quantify the frequency and magnitude of allele bias in a distinct way using clustering, after transforming the allele expression data for each gene in each tissue to a simpler format for clustering (See methods).

### Hierarchical Clustering by Allele Bias Across Tissues Identified Four Major Groups of Genes

We next used hierarchical clustering to cluster genes based on their allele bias (**Supplemental Figure 1**). The hierarchical clustering identified four major categories (clusters) of allele bias patterning (**Figure 2**). We found 4,792 biallelic genes, 3,539 slightly biased genes, 4,336 potentially very biased genes, and 653 mostly monoallelic genes. The heatmap does reveal some tissue-dependent differences where the degree of bias was a function of both gene and tissue (Sands, Yun, and Mendenhall 2021; Nag et al. 2015).

**Figure 2.**
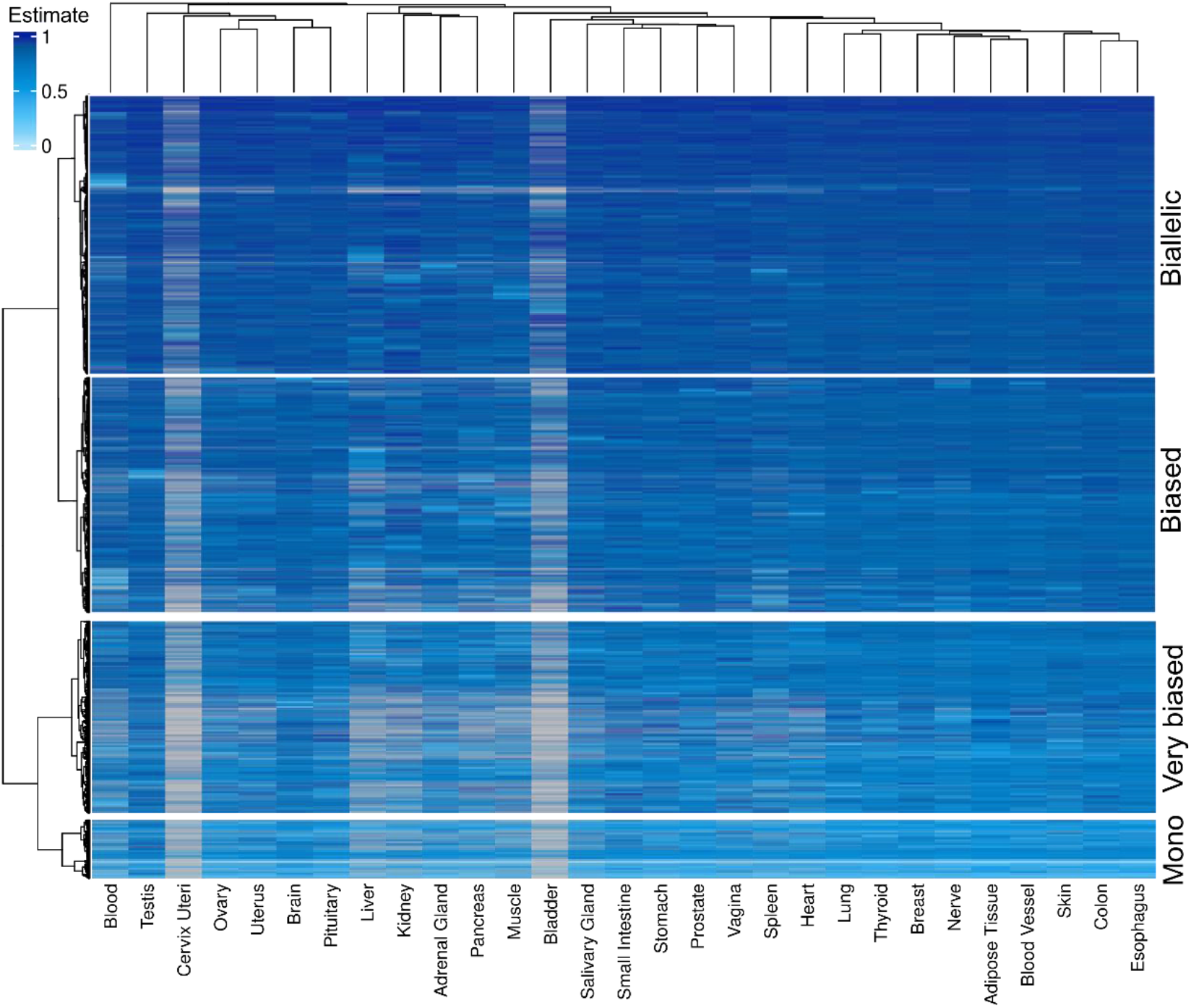
Genes classified by allele ratio estimate cluster into four distinct groups. Clustering algorithms identified four distinct classes of genes based on the transformed slope values for the data for each gene with discernible allele specific expression in all the tissues in which that gene was expressed above our threshold limits (See Methods).

We did not include known imprinted genes in our *main* analysis because they would skew the clustering of the random autosomal monoallelic expression. Imprinted genes always have a non-random parent of origin silencing in at least one tissue, which is usually critical for viability. Imprinting has been compared in more detail to random autosomal monoallelic expression previously (Chess 2016). Hierarchical clustering of the data showed three major groups when we included known imprinted genes listed in Tucci et al. (Tucci et al. 2019) (**Supplemental Figure 3**). Because some imprinted genes are only imprinted in one tissue and not others, some imprinted genes are classified as biallelic.

### Biased and Monoallelic Genes Affect Genetic Disease and Cancer

Genes associated with cancer and genetic disease are frequently expressed in a biased/monoallelic fashion, impacting the assumption of Mendelian biallelic expression. Both the silencing of individual genes and the many combinations of alleles silenced in individual bodies could account for incomplete penetrance and missing heritability for many traits and diseases.

**Figure 3A** shows that 29% of the 4,486 ORFs associated with human disease are expressed in a biased, very biased, or monoallelic fashion. For the purposes of our Venn Diagrams, we grouped biased and very biased together. Of the 4,486 genes on the OMIM list of human disease-related genes that we were able to classify, 1,509 are biallelic, 1,796 are biased, and 111 are monoallelic.. The degree of MAE observed in the GTEx data could be an underestimate (**Figure 3A)**; the GTEx data did not capture the clinically relevant silencing of *POUF1*, which we know can happen (Okamoto et al. 1994), but seemingly does not happen frequently among the hundreds of individuals sampled for the GTEx project.

**Figure 3.**
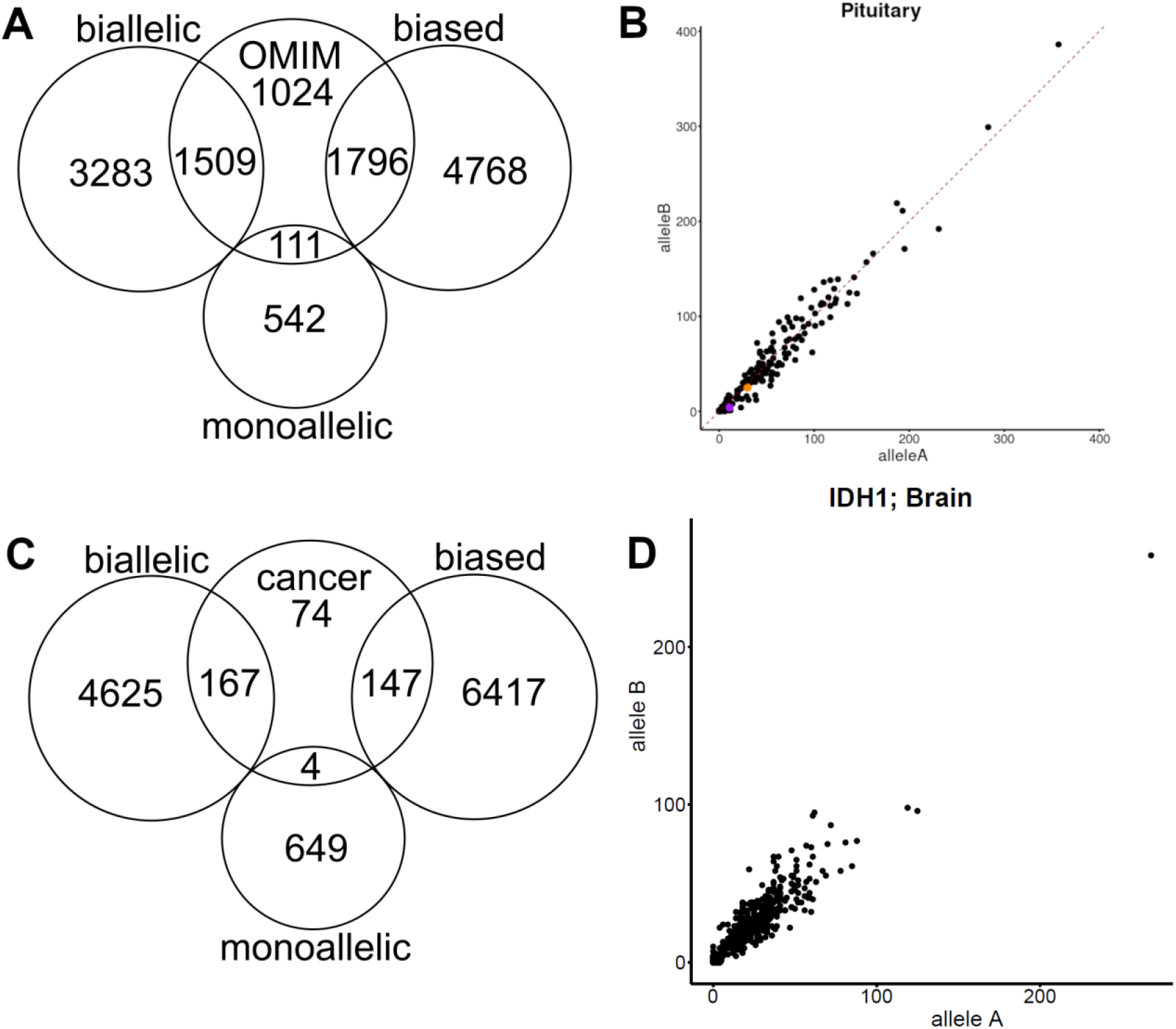
Biased and Monoallelically Expressed Genes are Enriched for Genes Affecting Genetic Disease and Cancer. **A)** Venn Diagrams of Genes classified as Biallelic, Biased/Very Biased, and Monoallelic by our classification scheme overlapping with gene associated with genetic disease. **B)** Scatter plot of *POUF1*/PIT1 expression in the pituitary gland. *POUF1*/PIT1 is known to be monoallelically expressed in some people in their pituitary glands (Okamoto et al. 1994), but was not captured by the GTEx data, indicating that the data is an underrepresentation of the possibilities for billions of individuals. The data is more likely a picture of what usually happens. **C)** Venn Diagrams of Genes classified as Biallelic, Biased/Very Biased, and Monoallelic by our classification scheme. **D)** Scatter plot of *IDH1* expression in the brain. *IDH1* is known to be monoallelically expressed in brain cancers (Walker et al 2012), but was not captured by the GTEx data, indicating that the data does not capture what goes on in cancer cells.

We next cross-referenced our gene clusters with 392 oncogenes and tumor suppressor genes as identified by Chakravarty et al. 2017. We found that 167 are biallelic, 147 are biased, and 4 are expressed in a monoallelic fashion (**Figure 3B**). Silencing of tumor suppressors and oncogenes can increase or decrease the risk of cancer, respectively. Silencing of a DNA repair gene like *BRCA1* in breast tissue can increase risk for cancer (X. Chen et al. 2008). Our approach did not classify *TP53* or *IDH1* as monoallelic or biased, but they have previously been observed exhibiting this behavior in brain tumors (Walker et al. 2012). A scatter plot of *IDH1* expression plotted by allele reads for each person’s brain tissue sample is shown in **Figure 3B, panel D**, revealing the expression of *IDH1* in normal tissues to be biallelic on average, with no incidences of major bias detected in these samples. *IDH1* is expressed monoallelically in about 25% of oligodendrogliomas, where it indicates differences in survival (Walker et al. 2012). See **Supplemental Figure 3** for the Venn diagram showing the overlap in classifications with the imprinted genes listed in Tucci et al. 2019 (Tucci et al. 2019).

### There are Tissues and Individuals with More Random Autosomal Allele Silencing

Given that there are more variably expressed genes, in terms of allele expression bias, we hypothesized that there may be tissues and individuals with relatively more allele bias. We quantified intrinsic noise (a quantitative measure of allele bias that measures deviation from a 1:1 ratio) for individual tissues and people and found that there were both noisier tissues and noisier individuals. We found that the testes and ovaries were the noisiest tissues, and that some people have much more random silencing of autosomal alleles than others. **Figure 4A** shows that tissues vary in the amount of intrinsic noise. We used the intrinsic noise metric for people and found that some people had far more silencing events than others, shown in **Figure 4B**, indicating that there may be a magnitude component to global silencing events in early development. This is consistent with the variation in silencing patterns in adult C. elegans resulting from variable activity of the histone H3K9 methyltransferase *SET-25/SUV39H1/EHMT2* in the early embryo (Sands et al. 2024).

**Figure 4:**
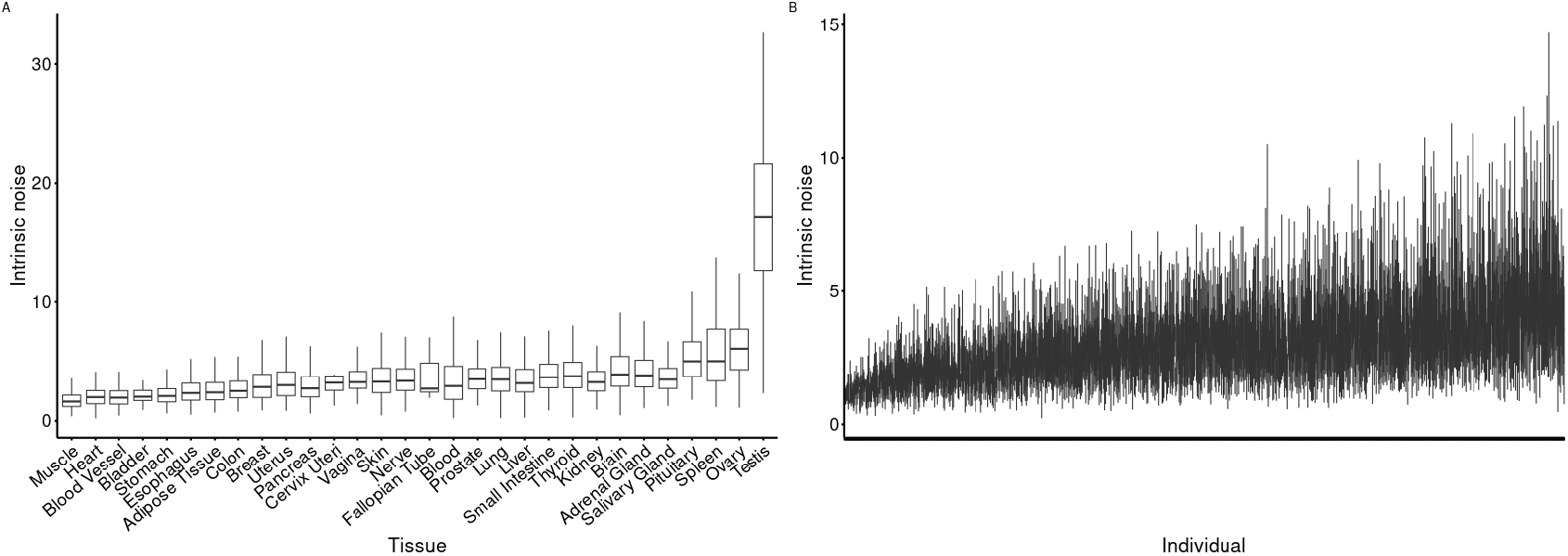
Some Tissues and Some Individuals have Relatively More Allele Expression Bias/Monoallelic Expression. **A)** Median intrinsic noise for each tissue; boxplots show mean and 25th, 75th percentiles of gene noise. Intrinsic noise is a measure of the deviation of allele expression from a 1:1 ratio. **B)** Intrinsic noise calculated for all genes in all tissues for each individual in the GTEx v8 dataset. There are clearly people with more and less biased/monoallelic expression. These genome-wide differences in silencing and bias may prove advantageous or disadvantageous in different scenarios.

### Age-related Changes in Autosomal Allele Silencing are Not the Same in All Tissues

There are changes in silencing with age, including at the level of individual alleles We determined if there were differences in which alleles are silenced in each tissue as a function of age. The strongest signal of allele ratio relationship change with age occurred in the brain (308 ORFs with FDR < 0.05), followed by blood vessel (11 ORFs with FDR < 0.05) (**Figure 5A, 5B**). We further explored allelic ratios in brain tissue. PRDX5, a gene known to aid in maintaining protein homeostasis and DNA damage while being linked to age-related diseases like Parkinson’s (Szeliga 2020; Agborbesong et al. 2023), changed from a biased allele ratio to more monoallelic with advancing age (FDR = 5.0 x 10^−27^, β = −3.3 x 10^−4^). We also found that UBB (the ubiquitin B gene) is monoallelically expressed, but trends toward biallelic with increased age (FDR = 1.3 x 10^−24^, β = 4.6 x 10^−2^). Ubiquitin, one of the most highly conserved proteins across the animal kingdom, plays crucial roles in the cell cycle and maintaining protein homeostasis by recruiting damaged or misfolded proteins to the proteasome. UBB has been repeatedly linked to diseases in the brain, such as Alzheimer’s (Wagh and Glickman 2025; Tan et al. 2007; Hope et al. 2003; Krishna-K, Behnisch, and Sajikumar 2022).

**Figure 5:**
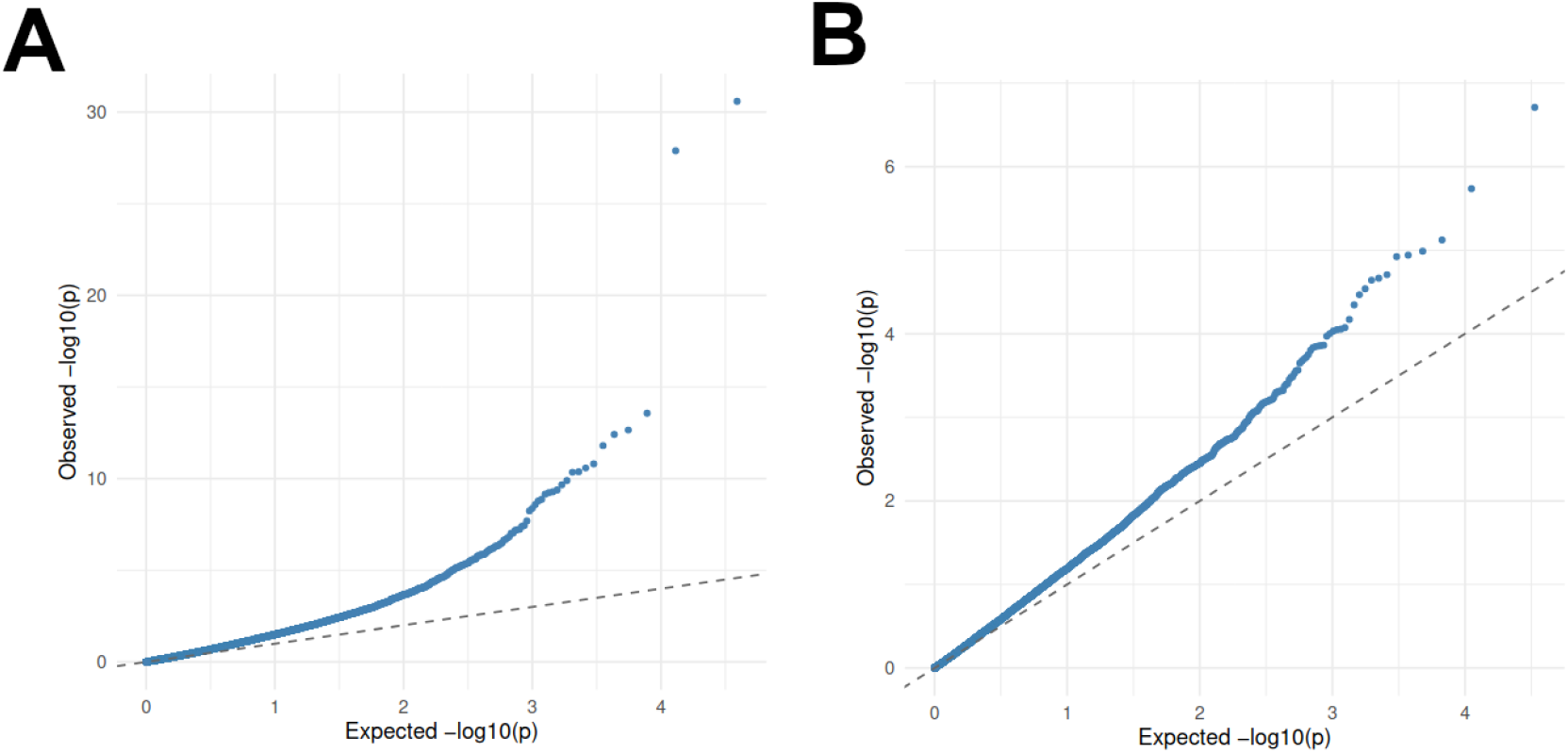
ORFs in the brain have great allele ratio changes with age. QQ plots depicting the signal of genes that change allele ratio with age in the (A) brain and (B) blood vessel.

### Individual People Maintain Unique Allele Expression Bias Combinations Across Tissues

In animal models, allelic imbalance in the few examined genes is sometimes propagated throughout tissues if silencing events happen early enough in development. We tested whether people maintained bias across tissues. If people were found to have the same bias for the same gene across different tissues, it would indicate that there was a silencing event early development before tissues differentiate and that silencing was maintained as tissues differentiated and mitotically grew.

Individuals in **Figure 6** are highlighted in different colors. The individuals are always seen on the same side of the 45-degree line in scatter plots of alleles of the same gene expressed in different tissues by different individuals. **Supplemental Figure 4** shows that when individual tissues were resequenced, they had the same or similar allele bias, indicating that these patterns are not created by stochastic sequencing noise. Individuals maintaining the same allele bias across tissues indicates that the medical cases in which buccal swabs were used to assess the state of expression in, for example, the pituitary gland, may have established a practice that can be maintained for the assessment of the silencing of many different genes in many different tissues, provided they are also expressed in the biopsied tissue. **Figure 6** also shows that different individuals have different combinations of allele bias. These different interindividual combinations of allele bias have tremendous potential to create diverse physiological states among individuals.

**Figure 6:**
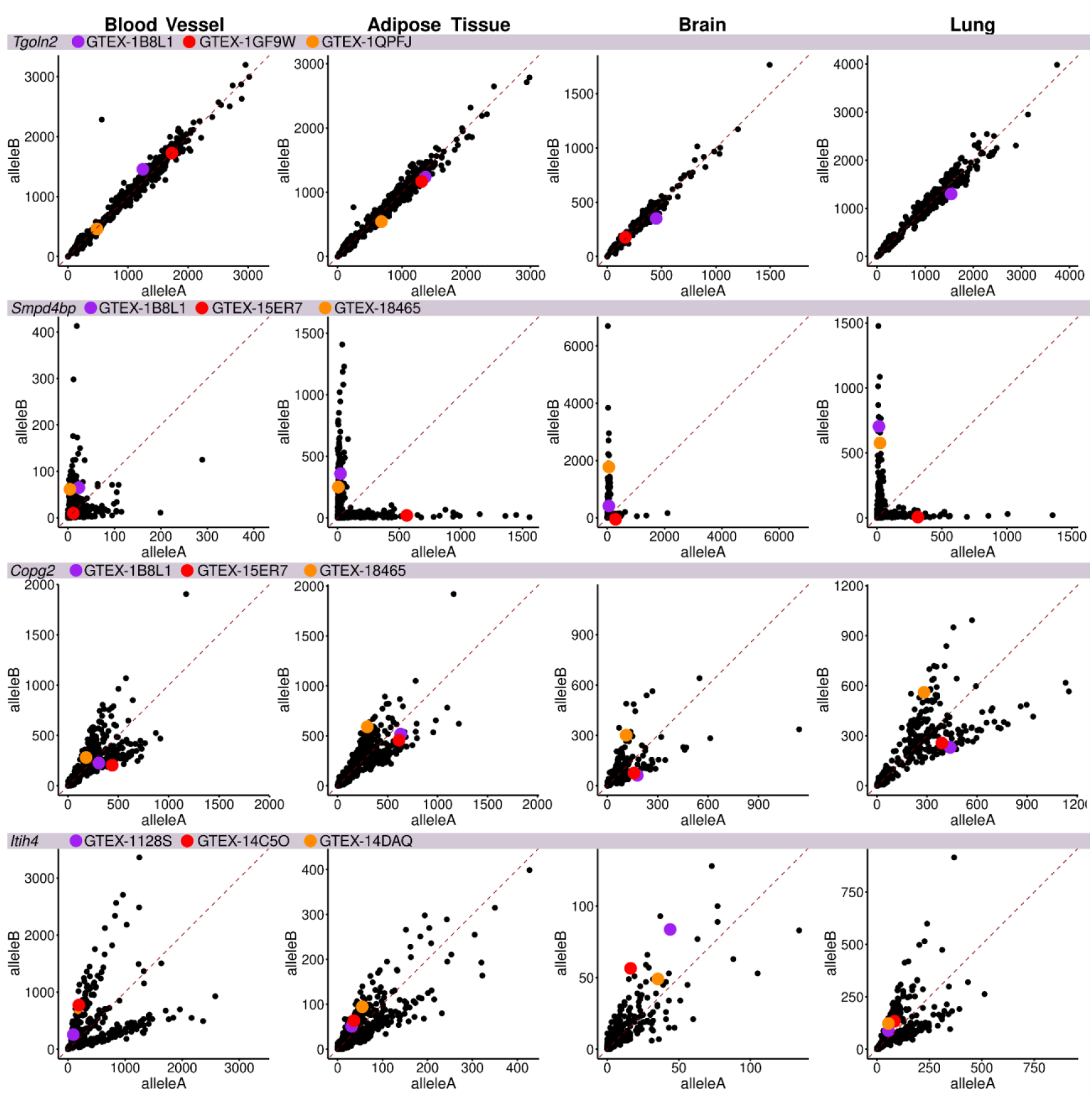
Individuals Maintain Distinct Combinations of Allele Expression Bias Across Tissues. Different individuals have different allele biases that are maintained across tissues that exhibit bias in gene expression. Variation in the degree of allele preference is widespread and can depend on tissue type. When there is bias, it is maintained across tissues for individuals, with three individuals highlighted with red, orange, and purple dots. Dots of the same color indicate the same individual. Row 1: *Tgoln2*, Row 2: *Copg2*. Row 3: S*mpd4bp*. Row 4: *Itih4*. The red dashed line is the x = y line.

## Discussion

We found that allele expression bias profiles are unique to each person and are likely established during early development because the bias of individual alleles is propagated across tissues. The propagation of bias across tissues allows us to infer bias about one tissue from sampling another. While the data reveal that some people maintain bias across tissues, the data do not exclude the possibility some people do not. We found there are both tissues that tend to have more allele bias and individuals that have more overall allele bias; these findings help us understand how random silencing can be more consequential/prevalent in particular organs and in particular individuals. Discrete traits, quantitative traits and risk of genetic disease are affected by allele dosages and activities, which may be controlled by these probability-driven autosomal silencing processes. Recessive traits can manifest despite individuals being heterozygous for the pathological allele. Silencing of dominant disease alleles can allow them to be transmitted without consequence to the carrier. In humans, in one case, a father’s pathological allele of *POUF1* was silenced, while his daughter’s was not. The daughter inherited the allele, but not the silenced state, and as a result suffered from *POUF1*-linked pituitary disease (Okamoto et al. 1994). Recent findings show that inborn errors of immunity are also caused by random autosomal monoallelic expression (Stewart et al. 2025). Many traits are likely to be influenced by this kind of silencing. The fact that different individuals had different combinations of which alleles were silenced suggests that complex differences in patterns of allele silencing are causing variation in biological system outputs.

Our analyses detected strong changes in allele bias with age in brains and blood vessels but no other tissues, indicating a strong change in silencing/desilencing with age in these tissues, relative to other tissues. Allele biases, or lack thereof, may be setting up different physiological states for aging and chronic disease, effectively serving as epigenetic determinants, just like genetic determinants (e.g., growth factor or insulin signaling genes) of risk. And, while medical reports indicate that allele silencing can last a lifetime in buccal epithelial cells (e.g., Okamoto et al. 1994), other tissues seem less stable for silencing.

Increasing evidence shows consistent transcriptional desilencing is associated with aging in many organisms (Sen et al. 2016; Wood et al. 2016). For instance, promoting heterochromatin formation increases lifespan in *Drosophila* (Jones et al. 2016). Growing data suggest that genetic de-silencing can lead to a more rapid age progression (H. Chen et al. 2016; De Cecco et al. 2013; Gorbunova et al. 2021). More rigorous analyses of age-related changes in endogenous and acquired silencing in humans will be relevant to the understanding of chronic diseases.

We found 151 of the 392 oncogenes and tumor suppressors are subject to probabilistic silencing, based on detecting allele bias or monoallelic expression for those genes in healthy tissues. Monoallelic expression is widely associated with many cancers, including leukemia, colorectal cancer, and breast cancer (Walker et al. 2012; Wei et al. 2013; Curia et al. 2012). Random silencing of pathological alleles could explain why some individuals who carry high-risk susceptibility alleles do not develop cancer or do so at a later age (Martincorena et al. 2018; Kratz et al. 2015). The differences in the expression of these oncogenes like *IDH1* or tumor suppressors like *TP53* could potentiate or suppress cancer risk and cancer progression in ways that traditional genetic testing could not predict (Walker et al 2012).

Allele-specific epigenetic medicine is developing, with six siRNA-based drugs already FDA-approved and in clinical use for silencing genes in the United States (Padda et al. 2025). Allele-specific silencing of pathological alleles is a logical next step in epigenetic medicine, which requires us to consider allele-specific expression profiles. The scatter plots showing that individuals have distinct allele expression states and combinations of those states throughout their bodies are suggest a need for scientists and physicians to consider MAE/allele-specific expression profiling in the application of these new therapeutics.

## Methods

### Scatter Plotting

The bulk RNAseq haplotype matrices made available by the GTEx Consortium v8 were used for all the analyses. This dataset contains 15,253 samples from 54 human tissues compiled from 838 human donors (THE GTEX CONSORTIUM 2020; Castel et al. 2020). These read-backed haplotype matrices were generated by running phASER v1.0.1 using the whole-genome sequencing data. Reads for “allele A” and reads for “allele B” were then separated into two different matrices. Data were plotted as scatter plots.

We note the RNA-seq approach discerns between tissues comprised of cells that are generally biallelic, monoallelic, or mostly biased for each allele, but would not be able to discern variegated allele expression patterns. Variegated means cells in a tissue autonomously expressing one or both alleles, creating a patchwork. When these cells are sequenced together in bulk sequencing, the number of mRNA transcripts from each allele from each cell is close to exactly the same, so it looks like biallelic expression in bulk sequencing results. We excluded genes that are known to be imprinted from our analyses (Tucci et al. 2019).

### Clustering

For clustering and heat map generation, since phASER cannot discern whether an allele is paternal or maternal (Allele1.A and Allele1.B are arbitrary titles for ORF variants in an individual and thus cannot be translated to mean the same thing between individuals), we transformed the more highly expressed allele for each ORF in each individual (across all their available samples) to be “allele A” and the more lowly expressed allele to be “allele B.” Next, with these transformed data, we estimated the allele A expression to allele B expression using linear modeling (linear model: allele A reads ∼ allele B reads) for each ORF in each tissue type.’

### Individual Human Noise Analysis

We quantified the intrinsic noise, a quantitative measure of the degree of allele bias (deviation from 1:1 ratio), for each individual person for all genes with distinct SNPs in alleles and plotted the intrinsic noise for each person as a line (box-like) plot. Intrinsic noise = (x - y)^2 / (2 * (<x> * <y>)) where x and y are each cell’s allele expression values and <x> and <y> are the average values for each allele’s expression in each individual, as done in (Sands, Yun, and Mendenhall 2021).

### Bias Across Tissues Analysis

We quantified the intrinsic noise for all genes expressed in each organ and plotted the median intrinsic noise values as box plots. Intrinsic noise = (x - y)^2 / (2 * (<x> * <y>)) where x and y are each cell’s allele expression values and <x> and <y> are the average values for each allele’s expression in each individual, as done in (Sands, Yun, and Mendenhall 2021).

### Age Analysis

We found age estimates for each gene in each tissue across individuals, adjusted for sex (linear model: allele ratio ∼ Age + Sex). The allele ratio for each gene was found in each individual by dividing the allele with the fewest reads by the allele with the most reads. For all analyses, we chose a modest filter of requiring 10 reads across 10 different individuals for each tissue, respectively.

## Statements and declarations

None. No competing interests.

## Sources of Funding

This research was funded by National Institutes of Health grant P20GM121176, National Institutes of Health grant R01AG07077601, and Longevity Impetus Grant to M.A.M. This research was funded in part through the NIH/NCI Cancer Center Support Grant P30 CA015704 and by NIH NIA Nathan Shock Center Grant P30AG013280. T.J. was supported by NIH/NCI T32CA009515. This research was supported in part through funding from Seattle Translational Tumor Research (STTR).

## Disclosures

The authors declare no competing interests.

## Author Contributions

B.L.M., B.S., A.R.M., and M.A.M. designed the study. B.L.M. executed the data analysis. A.R.M. and M.A.M. provided advisory support for the analyses. A.R.M. and B.L.M. generated the first draft. All authors helped interpret results in the context of their various research and/or clinical specialties and helped revise the manuscript.

## Acknowledgements

We would like to thank Alexander Gimelbrant and Asya Mendelovich for thoughtful discussions about the manuscript that improved the quality of the manuscript.

## Supplemental Information 1

Allele’s assignment to detect haplotype expression is computationally arbitrary. The assignment of “Allele A” or “Allele B” is consistent in individuals but is done without knowing which allele is dominant/recessive or maternal/paternal. Thus, we reasoned that the haplotype expression data ought to be transformed in a way where we can consistently detect the degree of allele bias. We determined that the allele with the most aggregate reads was the allele A across all available tissues for each individual. Interestingly, the allele with the most reads was generally consistent across their tissue types, including in re-sequenced samples.

After this simple transformation, we fit linear models to estimate the relationship of allele-specific expression for each ORF in each tissue type across individuals. We next used a heatmap with hierarchical clustering to visualize these calculated degrees of bias for ORFs with at least one significant relationship between allele reads from the linear model (p_adj_ > 0.05; reads allele A ∼ reads allele B) in any tissue. Encouragingly, the ORFs clustered clearly in alignment with their estimates. Notably, there are very limited instances when the estimate is above 1, indicative that the allele with the most reads in an individual is consistently the allele with the most reads in all tissues.

Specifically, many ORFs are consistently biallelic across all tissues, some ORFs have monoallelic expression patterns across all tissues, some ORFs have biased expression patterns across all tissues, and several ORFs have mixed monoallelic, biased, and biallelic expression depending on the tissue type. Monoallelic and biased expression patterns consistently patterned allele A more favorably, indicating the consistent tendency for allele A to be expressed more highly than allele B across tissue types in an individual. So, we clustered every allele ratio estimate for every gene with reads in at least 50% of all samples. Encouragingly, more closely related tissues clustered, like the brain and the pituitary as well as the ovary and uterus. These data show that many ORFs have consistent degrees of allele bias across individuals; however, several ORFs have differential allele bias expression tendencies dependent on tissue type. Of the 12,117 ORFs that passed our filtering (27,225 ORFs total provided in the GTEx haplotype matrix), 6,989 are consistently biallelic, 4,882 show some allele bias, and 246 are consistently monoallelic.

This study inquired into the broad classification of how ORF preference is expressed across individuals in tissue types. This study used a simple and conceptually easy-to-understand method for classifying ORFs’ tendencies in having allele expression bias. However, some ORFs follow a monoallelic expression in less than a handful of tissues, which the particular clustering method we used misses. We can see that some ORFs in our dataset have this pattern of, for example, a widely biallelic pattern with a few tissue-specific cases of biased patterns of our Biallelic and Slightly Biased clusters, that will show diversity like Figures 2 and 3)that our hierarchical clustering method misses. Even though these ORFs show bias in only a small subset of tissues, this may indicate physiological and biological relevance. Future studies will use machine learning/AI to classify all scatter plots of all ORF expression in all tissues (by using, for example, image classification with deep learning) to draw out these instances of rare allele preference confidently. Still, the method and results here present findings that have otherwise been missed entirely– that there is an entire class of ORFs that show consistent allele preference through many tissues– that ought to be understood.

**Supplemental Figure 1:**
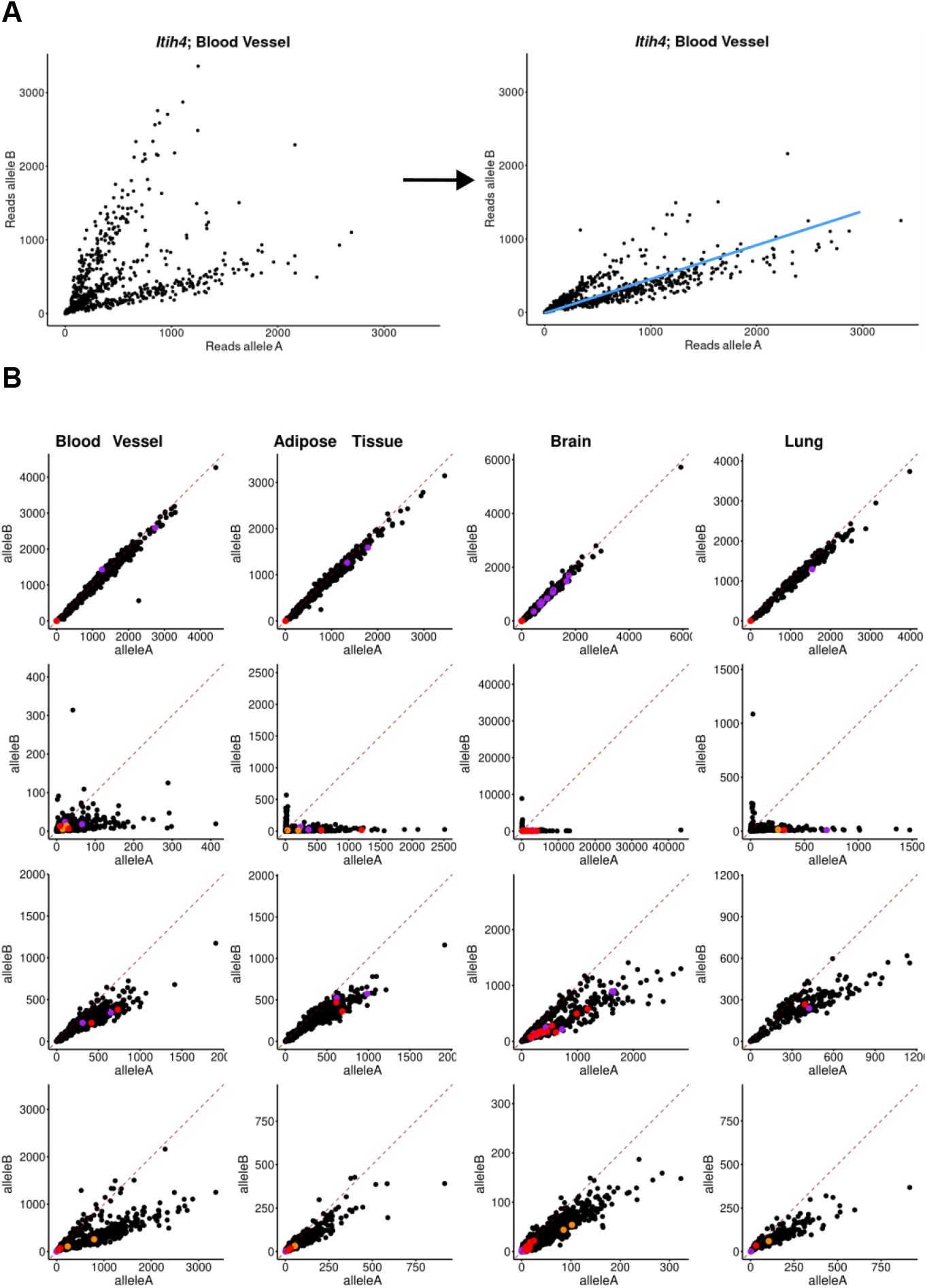
Transformed scatter plots. A) Blue line is a rough estimate of the linear model estimate of reads from allele A to B used for Figure 4 (linear model: allele A ∼ allele B). Estimates closer to 1 indicate a biallelic expression, whereas lower estimates indicate some allele bias. B) Dots of the same color indicate the same individual. Row 1: *Tgoln2*, Row 2: *Copg2*. Row 3: *Smpd4bp*. Row 4: *Itih4*. The red dashed line is the x = y line.

**Supplemental Figure 2.**
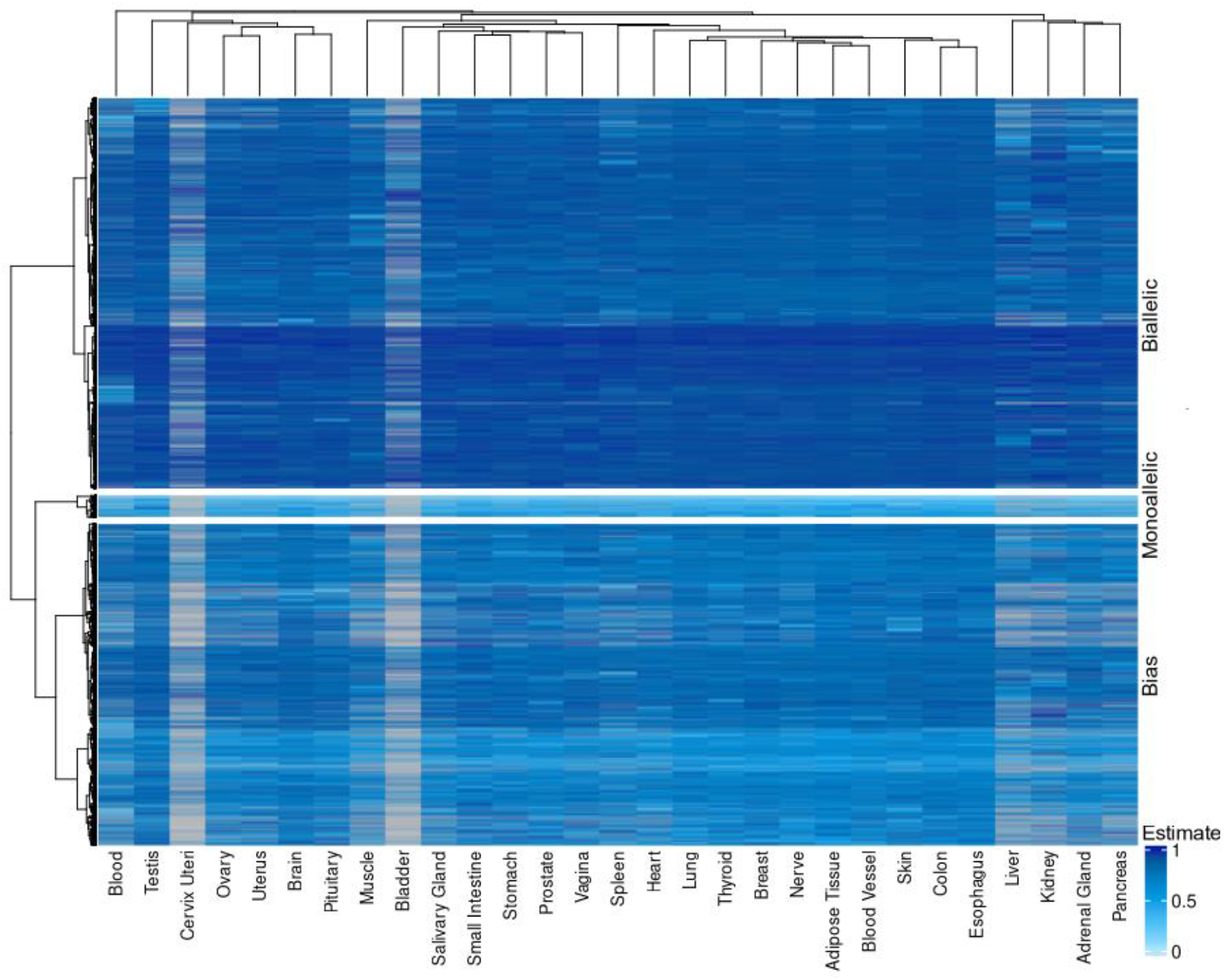
allele ratio estimate heatmap and clustering without filtering out imprinting genes.

**Supplemental Figure 3.**
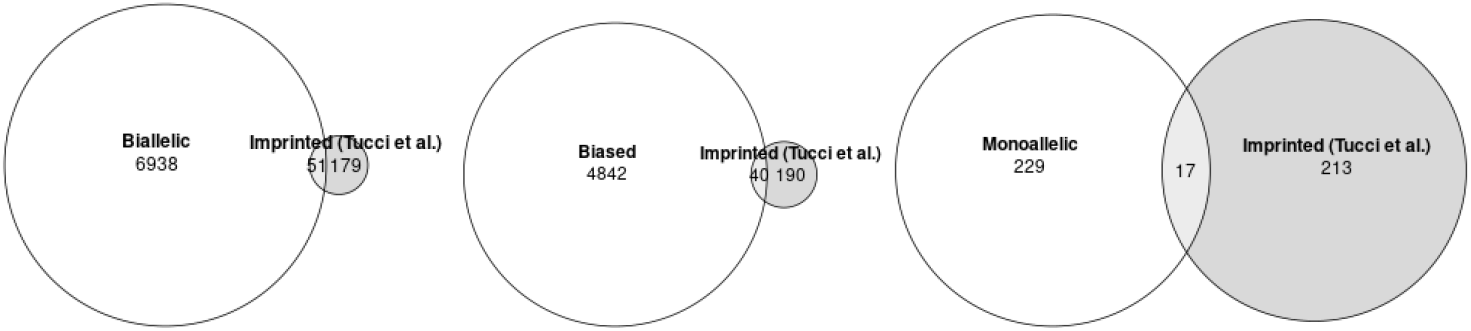
Venn diagrams of overlap our our findings with Tucci et al.

**Supplemental Figure 4:**
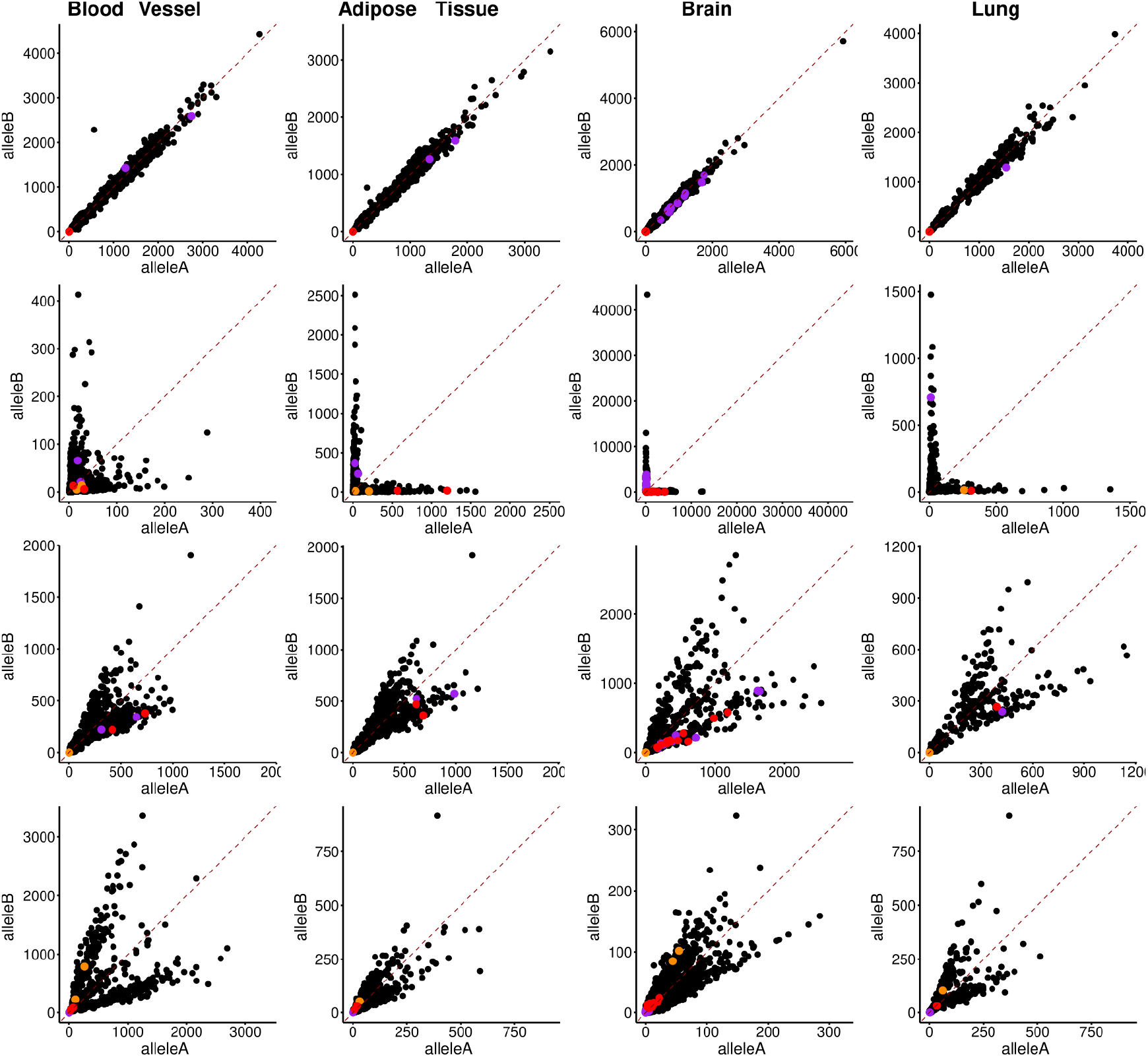
Resampling Shows the Same Allele Bias Direction. The resampling of tissues for RNA-seq revealed that the bias was not a stochastic artifact, as samples that were sequenced showed the same allele bias, just with more reads per sample.

## Notes

### Competing Interest Statement

The authors have declared no competing interest.

## References

Adegbola, Abidemi A., Gerald F. Cox, Elizabeth M. Bradshaw, David A. Hafler, Alexander Gimelbrant, and Andrew Chess. 2015. “Monoallelic Expression of the Human FOXP2 Speech Gene.” Proceedings of the National Academy of Sciences of the United States of America 112 (22): 6848–54.

Agborbesong, Ewud, Julie X. Zhou, Linda X. Li, Peter C. Harris, James P. Calvet, and Xiaogang Li. 2023. “Prdx5 Regulates DNA Damage Response through Autophagy-Dependent Sirt2-p53 Axis.” Human Molecular Genetics 32 (4): 567–79.

Antonarakis, S. E., S. H. Irkin, T. C. Cheng, A. F. Scott, J. P. Sexton, S. P. Trusko, S. Charache, and H. H. Kazazian Jr. 1984. “Beta-Thalassemia in American Blacks: Novel Mutations in the ‘TATA’ Box and an Acceptor Splice Site.” Proceedings of the National Academy of Sciences of the United States of America 81 (4): 1154–58.

Balliu, Brunilda, Matthew Durrant, Olivia de Goede, Nathan Abell, Xin Li, Boxiang Liu, Michael J. Gloudemans, et al. 2019. “Genetic Regulation of Gene Expression and Splicing during a 10-Year Period of Human Aging.” Genome Biology 20 (1): 230.

Bielski, Craig M., Mark T. A. Donoghue, Mayur Gadiya, Aphrothiti J. Hanrahan, Helen H. Won, Matthew T. Chang, Philip Jonsson, et al. 2018. “Widespread Selection for Oncogenic Mutant Allele Imbalance in Cancer.” Cancer Cell 34 (5): 852–62.e4.

Buckland, P. 2004. “Allele-Specific Gene Expression Differences in Humans.” Human Molecular Genetics 13 Spec No 2 (October):R255–60.

Burnaevskiy, Nikolay, Bryan Sands, Soo Yun, Patricia M. Tedesco, Thomas E. Johnson, Matt Kaeberlein, Roger Brent, and Alexander Mendenhall. 2019. “Chaperone Biomarkers of Lifespan and Penetrance Track the Dosages of Many Other Proteins.” Nature Communications 10 (1): 5725.

Castel, Stephane E., François Aguet, Pejman Mohammadi, François Aguet, Shankara Anand, Kristin G. Ardlie, Stacey Gabriel, et al. 2020. “A Vast Resource of Allelic Expression Data Spanning Human Tissues.” Genome Biology 21 (1): 234.

Castel, Stephane E., Alejandra Cervera, Pejman Mohammadi, François Aguet, Ferran Reverter, Aaron Wolman, Roderic Guigo, Ivan Iossifov, Ana Vasileva, and Tuuli Lappalainen. 2018. “Modified Penetrance of Coding Variants by Cis-Regulatory Variation Contributes to Disease Risk.” Nature Genetics 50 (9): 1327–34.

Chakravarty, Debyani, Jianjiong Gao, Sarah M. Phillips, Ritika Kundra, Hongxin Zhang, Jiaojiao Wang, Julia E. Rudolph, et al. 2017. “OncoKB: A Precision Oncology Knowledge Base.” JCO Precision Oncology 10.1200/PO.17.00011.

Chen, Haiyang, Xiaobin Zheng, Danqing Xiao, and Yixian Zheng. 2016. “Age-Associated de-Repression of Retrotransposons in the Drosophila Fat Body, Its Potential Cause and Consequence.” Aging Cell 15 (3): 542–52.

Chen, Xiaowei, Joellen Weaver, Betsy A. Bove, Lisa A. Vanderveer, Susan C. Weil, Alexander Miron, Mary B. Daly, and Andrew K. Godwin. 2008. “Allelic Imbalance in BRCA1 and BRCA2 Gene Expression Is Associated with an Increased Breast Cancer Risk.” Human Molecular Genetics 17 (9): 1336–48.

Chess, Andrew. 2016. “Monoallelic Gene Expression in Mammals.” Annual Review of Genetics 50 (1): 317–27.

Curia, Maria Cristina, Sabrina De Iure, Laura De Lellis, Serena Veschi, Sandra Mammarella, Marquitta J. White, Jacquelaine Bartlett, et al. 2012. “Increased Variance in Germline Allele-Specific Expression of APC Associates with Colorectal Cancer.” Gastroenterology 142 (1): 71–77.e1.

De Cecco, Marco, Steven W. Criscione, Edward J. Peckham, Sara Hillenmeyer, Eliza A. Hamm, Jayameenakshi Manivannan, Abigail L. Peterson, Jill A. Kreiling, Nicola Neretti, and John M. Sedivy. 2013. “Genomes of Replicatively Senescent Cells Undergo Global Epigenetic Changes Leading to Gene Silencing and Activation of Transposable Elements.” Aging Cell 12 (2): 247–56.

Eckersley-Maslin, Mélanie A., and David L. Spector. 2014. “Random Monoallelic Expression: Regulating Gene Expression One Allele at a Time.” Trends in Genetics: TIG 30 (6): 237– 44.

Eckersley-Maslin, Mélanie A., David Thybert, Jan H. Bergmann, John C. Marioni, Paul Flicek, and David L. Spector. 2014. “Random Monoallelic Gene Expression Increases upon Embryonic Stem Cell Differentiation.” Developmental Cell 28 (4): 351–65.

Falkenberg, Kim D., Nancy E. Braverman, Ann B. Moser, Steven J. Steinberg, Femke C. C. Klouwer, Agatha Schlüter, Montserrat Ruiz, et al. 2017. “Allelic Expression Imbalance Promoting a Mutant PEX6 Allele Causes Zellweger Spectrum Disorder.” The American Journal of Human Genetics 101 (6): 965–76.

Gandal, Michael J., Jillian R. Haney, Neelroop N. Parikshak, Virpi Leppa, Gokul Ramaswami, Chris Hartl, Andrew J. Schork, et al. 2018. “Shared Molecular Neuropathology across Major Psychiatric Disorders Parallels Polygenic Overlap.” Science 359 (6376): 693–97.

Gärtner, K. 1990. “A Third Component Causing Random Variability beside Environment and Genotype. A Reason for the Limited Success of a 30 Year Long Effort to Standardize Laboratory Animals?” Laboratory Animals 24 (1): 71–77.

Gärtner, Klaus. 2012. “Commentary: Random Variability of Quantitative Characteristics, an Intangible Epigenomic Product, Supporting Adaptation.” International Journal of Epidemiology 41 (2): 342–46.

Gendrel, Anne-Valerie, Mikael Attia, Chong-Jian Chen, Patricia Diabangouaya, Nicolas Servant, Emmanuel Barillot, and Edith Heard. 2014. “Developmental Dynamics and Disease Potential of Random Monoallelic Gene Expression.” Developmental Cell 28 (4): 366–80.

Gimelbrant, Alexander, John N. Hutchinson, Benjamin R. Thompson, and Andrew Chess. 2007. “Widespread Monoallelic Expression on Human Autosomes.” Science (New York, N.Y.) 318 (5853): 1136–40.

Gorbunova, Vera, Andrei Seluanov, Paolo Mita, Wilson McKerrow, David Fenyö, Jef D. Boeke, Sara B. Linker, et al. 2021. “The Role of Retrotransposable Elements in Ageing and Age-Associated Diseases.” Nature 596 (7870): 43–53.

Gupta, Saumya, Denis L. Lafontaine, Sebastien Vigneau, Asia Mendelevich, Svetlana Vinogradova, Kyomi J. Igarashi, Andrew Bortvin, Clara F. Alves-Pereira, Anwesha Nag, and Alexander A. Gimelbrant. 2022. “RNA Sequencing-Based Screen for Reactivation of Silenced Alleles of Autosomal Genes.” G3 (Bethesda, Md.) 12 (2): jkab428.

Hamosh, A., A. F. Scott, J. Amberger, D. Valle, and V. A. McKusick. 2000. “Online Mendelian Inheritance in Man (OMIM).” Human Mutation 15 (1): 57–61.

He, Liang, Yury Loika, and Alexander M. Kulminski. 2022. “Allele-Specific Analysis Reveals Exon- and Cell-Type-Specific Regulatory Effects of Alzheimer’s Disease-Associated Genetic Variants.” Translational Psychiatry 12 (1): 163.

Hope, Andrew D., Rohan de Silva, David F. Fischer, Elly M. Hol, Fred W. van Leeuwen, and Andrew J. Lees. 2003. “Alzheimer’s Associated Variant Ubiquitin Causes Inhibition of the 26S Proteasome and Chaperone Expression.” Journal of Neurochemistry 86 (2): 394–404.

Huang, F. W., C. M. Bielski, M. L. Rinne, W. C. Hahn, W. R. Sellers, F. Stegmeier, L. A. Garraway, and G. V. Kryukov. 2015. “TERT Promoter Mutations and Monoallelic Activation of TERT in Cancer.” Oncogenesis 4 (12): e176.

Izzi, Benedetta, Mariaelena Pistoni, Katrien Cludts, Pinar Akkor, Diether Lambrechts, Catherine Verfaillie, Peter Verhamme, Kathleen Freson, and Marc F. Hoylaerts. 2016. “Allele-Specific DNA Methylation Reinforces PEAR1 Enhancer Activity.” Blood 128 (7): 1003–12.

Jones, Brian C., Jason G. Wood, Chengyi Chang, Austin D. Tam, Michael J. Franklin, Emily R. Siegel, and Stephen L. Helfand. 2016. “A Somatic piRNA Pathway in the Drosophila Fat Body Ensures Metabolic Homeostasis and Normal Lifespan.” Nature Communications 7(December):13856.

Kimmel, Jacob C., Lolita Penland, Nimrod D. Rubinstein, David G. Hendrickson, David R. Kelley, and Adam Z. Rosenthal. 2019. “Murine Single-Cell RNA-Seq Reveals Cell-Identity- and Tissue-Specific Trajectories of Aging.” Genome Research 29 (12): 2088–2103.

Koivisto, U. M., J. J. Palvimo, O. A. Jänne, and K. Kontula. 1994. “A Single-Base Substitution in the Proximal Sp1 Site of the Human Low Density Lipoprotein Receptor Promoter as a Cause of Heterozygous Familial Hypercholesterolemia.” Proceedings of the National Academy of Sciences of the United States of America 91 (22): 10526–30.

Kratz, C. P., L. Franke, H. Peters, N. Kohlschmidt, B. Kazmierczak, U. Finckh, A. Bier, et al. 2015. “Cancer Spectrum and Frequency among Children with Noonan, Costello, and Cardio-Facio-Cutaneous Syndromes.” British Journal of Cancer 112 (8): 1392–97.

Kravitz, Stephanie N., Elliott Ferris, Michael I. Love, Alun Thomas, Aaron R. Quinlan, and Christopher Gregg. 2023. “Random Allelic Expression in the Adult Human Body.” Cell Reports 42 (1): 111945.

Krishna-K, Kumar, Thomas Behnisch, and Sreedharan Sajikumar. 2022. “Modulation of the Ubiquitin-Proteasome System Restores Plasticity in Hippocampal Pyramidal Neurons of the APP/PS1 Alzheimer’s Disease-like Mice.” Journal of Alzheimer’s Disease: JAD 86 (4): 1611–16.

Lee, Changhoon, Eun Yong Kang, Michael J. Gandal, Eleazar Eskin, and Daniel H. Geschwind. 2019. “Profiling Allele-Specific Gene Expression in Brains from Individuals with Autism Spectrum Disorder Reveals Preferential Minor Allele Usage.” Nature Neuroscience 22 (9): 1521–32.

Lee, Jun-Yeong, Ian Davis, Elliot H. H. Youth, Jonghwan Kim, Gary Churchill, James Godwin, Ron Korstanje, and Samuel Beck. 2021. “Misexpression of Genes Lacking CpG Islands Drives Degenerative Changes during Aging.” Science Advances 7 (51): eabj9111.

Luft, Juliet, Robert S. Young, Alison M. Meynert, and Martin S. Taylor. 2020. “Detecting Oncogenic Selection through Biased Allele Retention in The Cancer Genome Atlas.” bioRxiv. bioRxiv. 10.1101/2020.07.03.186593.

Martincorena, Iñigo, Joanna C. Fowler, Agnieszka Wabik, Andrew R. J. Lawson, Federico Abascal, Michael W. J. Hall, Alex Cagan, et al. 2018. “Somatic Mutant Clones Colonize the Human Esophagus with Age.” Science 362 (6417): 911–17.

McKean, David M., Jason Homsy, Hiroko Wakimoto, Neil Patel, Joshua Gorham, Steven R. DePalma, James S. Ware, et al. 2016. “Loss of RNA Expression and Allele-Specific Expression Associated with Congenital Heart Disease.” Nature Communications 7 (September):12824.

Nag, Anwesha, Virginia Savova, Ho-Lim Fung, Alexander Miron, Guo-Cheng Yuan, Kun Zhang, and Alexander A. Gimelbrant. 2013. “Chromatin Signature of Widespread Monoallelic Expression.” eLife 2 (December):e01256.

Nag, Anwesha, Sébastien Vigneau, Virginia Savova, Lillian M. Zwemer, and Alexander A. Gimelbrant. 2015. “Chromatin Signature Identifies Monoallelic Gene Expression Across Mammalian Cell Types.” 5 (8): 1713–20.

Okamoto, N., Y. Wada, S. Ida, R. Koga, K. Ozono, H. Chiyo, A. Hayashi, and K. Tatsumi. 1994. “Monoallelic Expression of Normal mRNA in the PIT1 Mutation Heterozygotes with Normal Phenotype and Biallelic Expression in the Abnormal Phenotype.” Human Molecular Genetics 3 (9): 1565–68.

Padda, Inderbir S., Arun U. Mahtani, Preeti Patel, and Mayur Parmar. 2025. “Small Interfering RNA (siRNA) Therapy.” In StatPearls. Treasure Island (FL): StatPearls Publishing.

Prashant, N. M., Hongyu Liu, Pavlos Bousounis, Liam Spurr, Nawaf Alomran, Helen Ibeawuchi, Justin Sein, Dacian Reece-Stremtan, and Anelia Horvath. 2019. “Estimating Allele-Specific Expression of SNVs from 10x Genomics Single-Cell RNA-Sequencing Data.” bioRxiv. bioRxiv. 10.1101/2019.12.22.886119.

Reinius, Björn, Jeff E. Mold, Daniel Ramsköld, Qiaolin Deng, Per Johnsson, Jakob Michaëlsson, Jonas Frisén, and Rickard Sandberg. 2016. “Analysis of Allelic Expression Patterns in Clonal Somatic Cells by Single-Cell RNA-Seq.” Nature Genetics 48 (11): 1430–35.

Sands, Bryan, Soo Yun, and Alexander R. Mendenhall. 2021. “Introns Control Stochastic Allele Expression Bias.” Nature Communications 12 (1): 6527.

Sands, Bryan, Soo R. Yun, Junko Oshima, and Alexander R. Mendenhall. 2024. “Maternal Histone Methyltransferases Antagonistically Regulate Monoallelic Expression in C. elegans.” bioRxiv.org: The Preprint Server for Biology, January, 2024.01.22.576748.

Savova, Virginia, Sung Chun, Mashaal Sohail, Ruth B. McCole, Robert Witwicki, Lisa Gai, Tobias L. Lenz, C-Ting Wu, Shamil R. Sunyaev, and Alexander A. Gimelbrant. 2016. “Genes with Monoallelic Expression Contribute Disproportionately to Genetic Diversity in Humans.” Nature Genetics 48 (3): 231–37.

Sen, Payel, Parisha P. Shah, Raffaella Nativio, and Shelley L. Berger. 2016. “Epigenetic Mechanisms of Longevity and Aging.” Cell 166 (4): 822–39.

Solovieff, Nadia, Chris Cotsapas, Phil H. Lee, Shaun M. Purcell, and Jordan W. Smoller. 2013. “Pleiotropy in Complex Traits: Challenges and Strategies.” Nature Reviews. Genetics 14 (7): 483–95.

Stewart, O’jay, Conor Gruber, Haley E. Randolph, Roosheel Patel, Meredith Ramba, Enrica Calzoni, Lei Haley Huang, et al. 2025. “Monoallelic Expression Can Govern Penetrance of Inborn Errors of Immunity.” Nature, January. 10.1038/s41586-024-08346-4.

Swisher, Elizabeth M., Kevin K. Lin, Amit M. Oza, Clare L. Scott, Heidi Giordano, James Sun, Gottfried E. Konecny, et al. 2017. “Rucaparib in Relapsed, Platinum-Sensitive High-Grade Ovarian Carcinoma (ARIEL2 Part 1): An International, Multicentre, Open-Label, Phase 2 Trial.” The Lancet Oncology 18 (1): 75–87.

Szeliga, M. 2020. “Peroxiredoxins in Neurodegenerative Diseases.” Antioxidants (Basel, Switzerland) 9 (November). 10.3390/antiox9121203.

Tan, Z., X. Sun, F-S Hou, H-W Oh, L. G. W. Hilgenberg, E. M. Hol, F. W. van Leeuwen, M. A. Smith, D. K. O’Dowd, and S. S. Schreiber. 2007. “Mutant Ubiquitin Found in Alzheimer’s Disease Causes Neuritic Beading of Mitochondria in Association with Neuronal Degeneration.” Cell Death and Differentiation 14 (10): 1721–32.

THE BRAINSTORM CONSORTIUM, Verneri Anttila, Brendan Bulik-Sullivan, Hilary K. Finucane, Raymond K. Walters, Jose Bras, Laramie Duncan, et al. 2018. “Analysis of Shared Heritability in Common Disorders of the Brain.” Science 360 (6395): eaap8757.

THE GTEX CONSORTIUM. 2020. “The GTEx Consortium Atlas of Genetic Regulatory Effects across Human Tissues.” Science 369 (6509): 1318–30.

Tucci, Valter, Anthony R. Isles, Gavin Kelsey, Anne C. Ferguson-Smith, Valter Tucci, Marisa S. Bartolomei, Nissim Benvenisty, et al. 2019. “Genomic Imprinting and Physiological Processes in Mammals.” Cell 176 (5): 952–65.

Vigneau, Sébastien, Svetlana Vinogradova, Virginia Savova, and Alexander Gimelbrant. 2018. “High Prevalence of Clonal Monoallelic Expression.” Nature Genetics 50 (9): 1198–99.

Wagh, Ajay R., and Michael H. Glickman. 2025. “Molecular Mechanisms Underlying p62-Dependent Secretion of the Alzheimer-Associated Ubiquitin Variant, UBB+1.” bioRxiv. 10.1101/2024.12.31.630908.

Walker, Erin J., Cindy Zhang, Pedro Castelo-Branco, Cynthia Hawkins, Wes Wilson, Nataliya Zhukova, Noa Alon, et al. 2012. “Monoallelic Expression Determines Oncogenic Progression and Outcome in Benign and Malignant Brain Tumors.” Cancer Research 72 (3): 636–44.

Wei, Quan-Xiang, Rainer Claus, Thomas Hielscher, Daniel Mertens, Aparna Raval, Christopher C. Oakes, Stephan M. Tanner, et al. 2013. “Germline Allele-Specific Expression of DAPK1 in Chronic Lymphocytic Leukemia.” PloS One 8 (1): e55261.

Wood, Jason G., Brian C. Jones, Nan Jiang, Chengyi Chang, Suzanne Hosier, Priyan Wickremesinghe, Meyrolin Garcia, et al. 2016. “Chromatin-Modifying Genetic Interventions Suppress Age-Associated Transposable Element Activation and Extend Life Span in Drosophila.” Proceedings of the National Academy of Sciences of the United States of America 113 (40): 11277–82.

Xu, Jin, Ava C. Carter, Anne-Valerie Gendrel, Mikael Attia, Joshua Loftus, William J. Greenleaf, Robert Tibshirani, Edith Heard, and Howard Y. Chang. 2017. “Landscape of Monoallelic DNA Accessibility in Mouse Embryonic Stem Cells and Neural Progenitor Cells.” Nature Genetics 49 (3): 377–86.

Yan, Hai, Weishi Yuan, Victor E. Velculescu, Bert Vogelstein, and Kenneth W. Kinzler. 2002. “Allelic Variation in Human Gene Expression.” Science (New York, N.Y.) 297 (5584): 1143.

